# Changes in the microbiome of the trophosome of *Lamellibrachia satsuma* induced by rearing

**DOI:** 10.64898/2026.04.29.721791

**Authors:** Tomoko Koito, Maui Tahara, Ryota Taira, Ayuta Yamaki, Makoto Sugimura, Hiroko Makita, Tomoko Yamamoto, Toshiro Yamanaka

## Abstract

**Background:** Adult vestimentiferan tubeworms inhabiting hydrothermal vents and cold seeps lack a mouth and anus and rely entirely on organic matter produced by sulfur -oxidizing autotrophic bacterial symbionts in their trophosomes. These symbionts, which predominantly belong to the genus *Proteobacteria*, are acquired horizontally from the environment. However, the effects of rearing conditions that differ from natural habitats on the microbiome composition or abundance of these bacteria remain unclear.

**Methods:** We conducted a metagenomic analysis of *Lamellibrachia satsuma* reared in an aquarium under sulfide-supplemented and sulfide-free conditions.

**Results:** Immediately after collection, the microbiome was dominated by known symbionts within γ-Proteobacteria, exhibiting low species diversity. After 6 months of rearing, the abundance of these symbionts significantly decreased under both conditions, whereas overall bacterial diversity increased. In particular, α-Proteobacteria became more abundant under sulfide-supplemented conditions, while δ-Proteobacteria predominated in the absence of sulfide. Despite these changes, symbionts were not entirely lost, and the hosts survived for 6 months, likely due to their low metabolic rate. These findings suggest that the microbiome of *L. satsuma* can respond flexibly to changes in the rearing environment. They also indicate that the host’s metabolism can be maintained even with a smaller quantity of symbiotic bacteria.

## Introduction

Vestimentiferan tubeworms, which are annelids that inhabit hydrothermal vents, methane seeps, and whale carcasses, are important members of chemosynthetic ecosystems (Black et al., 1997). Adult tubeworms exhibit a unique ecology in which the mouth and anus disappear during growth (Jones, 1981; Bright; Klose & Nussbaumer, 2013). Trophosomes are the primary organ in the body.

Many symbiotic bacteria use environmental hydrogen sulfide as a substrate to fix carbon through the Calvin–Benson–Bassham pathway and the reductive tricarboxylic acid cycle (Cavanaugh et al., 1981; Jones, 1981; Bright; Klose & Nussbaumer, 2013; Patra; Kwon & Yang, 2022). Thus, instead of feeding independently, adult tubeworms rely on organic matter produced by their symbionts for sustenance (Felbeck & Childress, 1988; Yorisue et al., 2012).

Tubeworms have been suggested to be transmitted horizontally through the uptake of symbionts from the environment (Jones & Gardiner, 1988). Similar to other polychaetes, tubeworm larvae possess a mouth and anus and grow during feeding. During development, free-living bacteria from the environment infect the larvae, thereby initiating symbiosis, promoting trophosome formation, and eventually leading to settlement (Gibson et al., 2010). Indeed, analysis of the tubeworm microbiome using a clone library method detected γ-Proteobacteria and ε-Proteobacteria, which were shown to be genetically related to species detected in the surrounding sediment (Patra et al., 2016). Free-living symbiotic bacteria have also been identified in biofilms and ambient water surrounding tubeworm colonies (Harmer et al., 2008).

Several reports have documented multiple symbioses in tubeworms, such as the detection of α, β, and γ-Proteobacteria by 16S rRNA analysis of the trophosome of *Lamellibrachia* sp. inhabiting off Hatsushima in Sagami Bay (Kimura; Higashide & Naganuma, 2003). Another case has been reported for *Lamellibrachia anaximandri* (Zimmermann et al., 2014). Although the functions and interactions of these bacteria within tubeworms remain unclear, the dominant symbiont species differ according to the habitat and species (Naganuma et al., 2005).

To survive, tubeworms require a consistent supply of sufficient substrate, such as dissolved sulfide, to support the growth of their symbiotic bacteria. In contrast, aquarium conditions differ from natural environment conditions. *Lamellibrachia satsuma*, which inhabits Kagoshima Bay, the shallowest habitat in the world, has been studied in aquariums (Kojima, 2002; Miyake et al., 2006). Their survival was achieved through addition of a sodium sulfide solution to the exhibition tank. In water, sodium sulfide solutions dissociates into H_2_S and HS^-^. Therefore, sodium sulfide has been widely used as an alternative to hydrogen sulfide addition for the rearing of animals that live in deep-sea hydrothermal vents and cold seeps and require hydrogen sulfide as a substrate for symbiotic bacteria (Goffredi et al., 1997; Kádár et al., 2005; Koito et al., 2010). Although sodium sulfide solution is used as a substitute for gaseous hydrogen sulfide in the maintenance of organisms constituting chemosynthetic ecosystems, whether this approach maintains the microbiome balance within the trophosome remains unclear. In addition, whether the microbiome can be maintained in sulfide-free culture conditions remains unknown.

In this study, we compared the microbiomes of *L. satsuma* immediately after collection with those of individuals reared under sulfide-supplemented or non-supplemented conditions. Although tubeworms have been reported to take up sulfide from sediment (Julian et al., 1999; Mizota & Yamanaka, 2003), sulfide was added to the rearing water in this study because adding an accurate amount of sulfide to the sediment is difficult.

## Materials & Methods

### Sample collection

*L. satsuma* was collected using a grabber from Kagoshima Bay in June 2018 on the 64th cruise of the Kagoshima University training ship *Nansei-maru*. They were transported to the Enoshima Aquarium, where six were immediately dissected (Initial). The remaining individuals were divided into two groups, with 12 individuals each: sulfide addition (hereafter, with-S group) and non-addition (hereafter, without-S group) tanks. Both experimental groups were maintained in a circulating filtration tank with a water volume of approximately 120 L (W750 mm × D400 mm × H500 mm). The rearing water consisted of natural seawater maintained at 15°C, which was supplied and circulated at random intervals to maintain water quality and adjust pH. In the with-S group, 50 mL of a 20% sodium sulfide solution was added to the sulfide addition tank every 1 h over 2 min, and the total sulfide concentration was adjusted to approximately 50 μg/L. Sulfide concentrations during the rearing period were measured 10 min after sulfide addition using a HACH® apparatus and reagent (TNT861). However, one individual from the without-S group died during the experiment. Therefore, six individuals from the with-S group and five from the without-S group were included in this study. After 6 months of rearing, samples were collected from each experimental group and dissected. The trophosomes were removed from the tubes, excised using sterilized scissors into 1.5 mL tubes, frozen in liquid nitrogen, and stored at - 80°C until use.

### Metagenomic analyses

DNA was extracted from the trophosomes of *L. satsuma*, following a previously reported method (Koito et al., 2023). The DNA extracts were sent to Bioengineering Lab. Co., Ltd. (Kanagawa, Japan) for metagenomic analysis. The concentrations of DNA solutions were measured using SynergyLX (BioTek) and Quanti Fluords DNA System (Promega). Library preparation was performed using two-step tailed PCR. The concentrations of the prepared libraries were measured using SynergyH1 (BioTek) and the Fluords DNA System. The generated libraries were quality checked using a Fragment Analyzer and dsDNA 915 Reagent Kit (Advanced Analytical Technologies) and sequenced at 2 × 300 bp using the MiSeq system and MiSeq Reagent Kit 3 (Illumina). The Fastq_barcode_ splitter of Fastx toolkit was used to extract only reads whose sequence start perfectly matched the primer used. Primer sequences were removed from the extracted sequences. Sequences with a quality score below 20 were removed using sickletools, and those shorter than 130 bases as well as paired sequences were discarded. The paired-end merge script FLASH was used to combine reads that passed the quality filtering. The merging parameters were as follows: merged sequence length, 260 bases; base length of the read, 230 bases; and minimum overlap, 10 bases. Merged sequences were subsequently filtered by base length, and only those between 246–260 bases were used for further analyses. The binding sequences that passed the filtering were removed using the UCHIME algorithm. The database consisted of 97% of the operational taxonomic units (OTUs) from SILVA (ver. 132). The non-chimeric sequences were extracted for further analysis. OTUs were generated using QIIME workflow scripts without reference sequences and with all parameters set to default conditions.

### Microbiome analyses

OTUs were aggregated at the phylum level, and the mean and standard deviation were calculated for each group. The counts for each OTU were aggregated, and Shannon diversity indices were calculated using a free program available on the website of the Japan Sea National Fisheries Research Institute (http://jsnfri.fra.affrc.go.jp/gunshu/tayodo.html). Multiple comparisons were performed using Tukey’s test. Linear discriminant analysis (LDA) effect size (LEfSe; http://huttenhower.sph.harvard.edu/galaxy/) analysis was used to identify differential abundances at the class level. To examine the sharing and specificity of OTUs across experimental groups, Venn diagrams were generated using a free web-based program (http://bioinformatics.psb.ugent.be/cgi-bin/liste/Venn/calculate_venn.htpl).

### Phylogenetic analysis

A phylogenetic tree was constructed using the maximum-likelihood method based on the top 10 OTUs with the highest proportion of total counts in each experimental group, along with sequences of *Lamellibrachia* symbionts obtained from DDBJ/EMBL/GenBank in previous studies. Optimal analytical conditions were determined using the ‘Find Best DNA/Protein Models (ML)’ function implemented in the free analysis software MEGA version 10.0.5. Phylogenetic analysis was performed with 1,000 bootstrap replications using the General Time Reversible model, which showed the highest likelihood. In this analysis, the rate was set as gamma-distributed.

### Quantitative real-time PCR

Quantitative real-time PCR was performed for Denovo1132, which had the highest abundance ratio across all experimental groups, to compare its abundance between groups. Primers for the dominant symbiont belonging to γ-Proteobacteria of *L. satsuma* were designed based on the Denovo1132 sequence. The primers used were as follows: GAM16SatsuF1, AAGTGGAATTCCGGGTGTAGC, GAM16SatsuR, and TAAGGTTCTTCGCGTTGCATC. The total DNA concentrations for the initial, with-S, and without-S groups were adjusted to 30 ng/µL using NanoDrop Lite. Each quantitative PCR (10 µL total) contained 5 µL of TB green, 0.3 μL of forward primer, 0.3 μL of reverse primer, and 1 μL of template. PCR cycling conditions were as follows: 95℃ for 30 s, followed by 50 cycles of 95℃ for 5 s, 55℃ for 30 s, and 72℃ for 30 s. A melting curve analysis was performed for each PCR amplicon.

Data are presented as the mean ± SE of relative DNA levels. The mean 16S rDNA level in the initial group was set to 1. Statistical analyses were performed using analysis of variance (ANOVA) followed by Tukey-Kramer post hoc tests.

## Results

### Microbiome analyses

The Shannon index increased in the following order: Initial, without-S, with-S, and without-S (Fig. 1). The mean values were 1.040, 1.422, and 2.307, respectively (Fig. 1). However, the differences were not statistically significant (ANOVA; Tukey-Kramer post hoc test, *p*=0.1524).

**Figure 1.**
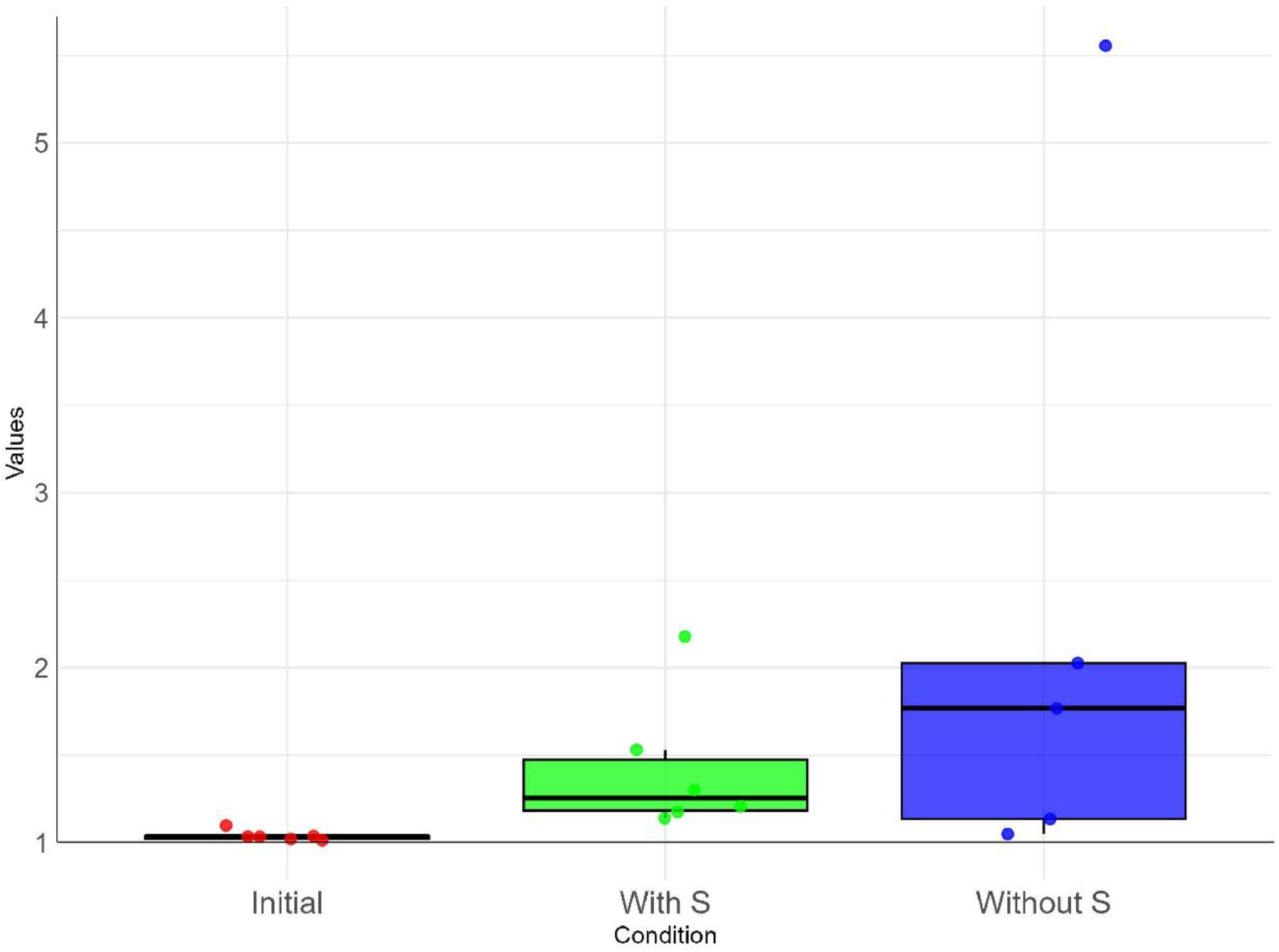
Shannon α-diversity index at the OTU level. No significant differences in microbial richness were observed among the three experimental groups (Tukey-Kramer; *p* > 0.05).

Phylum-level analysis revealed the presence of 1 archaeal phylum and 23 bacterial phyla across the experimental group (Table 1). No archaea were detected in the control group. Thirteen phyla were detected in the initial group, whereas 9 and 14 phyla were detected in the with-S and without-S groups, respectively. Bacteroidetes, Epsilonbacteraeota, Firmicutes, and Proteobacteria were detected in all experimental groups (Table 1).

**Table 1.** Phylum-level composition of the bacterial flora in the Initial, with-S, and without-S trophosomes of *Lamellibrachia satsuma*.

| Kingdom | Phyla | Initial | se | With sulfide | se | Without sulfide | se |
| --- | --- | --- | --- | --- | --- | --- | --- |
| Archaea | Thaumarchaeota | 0 | 0 | 0.001 | 0.001 | 0.242 | 0.242 |
| Bacteria | Acidobacteria | 0.011 | 0.011 | 0 | 0 | 0 | 0 |
|  | Actinobacteria | 0.004 | 0.003 | 0 | 0 | 0 | 0 |
|  | Aegiribacteria | 0.007 | 0.006 | 0 | 0 | 0 | 0 |
|  | Bacteroidetes | 0.006 | 0.003 | 1.210 | 0.609 | 1.420 | 1.168 |
|  | Calditrichaeota | 0.002 | 0.002 | 0 | 0 | 0 | 0 |
|  | Chloroflexi | 0.006 | 0.006 | 0 | 0 | 0 | 0 |
|  | Cloacimonetes | 0 | 0 | 0 | 0 | 0.011 | 0.011 |
|  | Cyanobacteria | 0 | 0 | 0.001 | 0.001 | 0.029 | 0.029 |
|  | Dadabacteria | 0 | 0 | 0 | 0 | 0.017 | 0.018 |
|  | Deferribacteres | 0 | 0 | 0 | 0 | 0.006 | 0.006 |
|  | Deinococcus-Thermus | 0 | 0 | 0 | 0 | 0.005 | 0.005 |
|  | Dependentiae | 0.001 | 0.001 | 0 | 0 | 0 | 0 |
|  | Epsilonbacteraeota | 0.002 | 0.002 | 0.126 | 0.091 | 0.478 | 0.352 |
|  | Firmicutes | 0.009 | 0.005 | 0.059 | 0.037 | 3.309 | 2.825 |
|  | Fusobacteria | 0.001 | 0.001 | 0 | 0 | 0.295 | 0.295 |
|  | Lentisphaerae | 0 | 0 | 0 | 0 | 0.041 | 0.373 |
|  | Modulibacteria | 0.005 | 0.005 | 0 | 0 | 0 | 0 |
|  | Nitrospirae | 0 | 0 | 0.005 | 0.005 | 0 | 0 |
|  | Patescibacteria | 0.009 | 0.009 | 0 | 0 | 0 | 0 |
|  | Planctomycetes | 0 | 0 | 0.091 | 0.075 | 0.044 | 0.043 |
|  | Proteobacteria | 99.934 | 0.047 | 98.491 | 0.741 | 94.085 | 4.243 |
|  | Spirochaetes | 0 | 0 | 0 | 0 | 0.004 | 0.004 |
|  | Verrucomicrobia | 0 | 0 | 0.004 | 0.004 | 0 | 0 |
| unclassified |  | 0.003 | 0.003 | 0.010 | 0.008 | 0.013 | 0.008 |

Intergroup comparisons revealed that Chromatiales (γ-Proteobacteria) exhibited a significantly higher abundance ratio in the initial group (Fig. 2). The abundance ratios of Kordiimonadates and Rhodobacterales (α-Proteobacteria), Thiotrichales (γ-Proteobacteria), and Flavobacteriales (Bacteroidetes) were significantly higher in the with-S group, whereas that of δ-Proteobacteria (δ-Proteobacteria) was significantly higher in the without-S group (Fig. 2).

**Figure 2.**
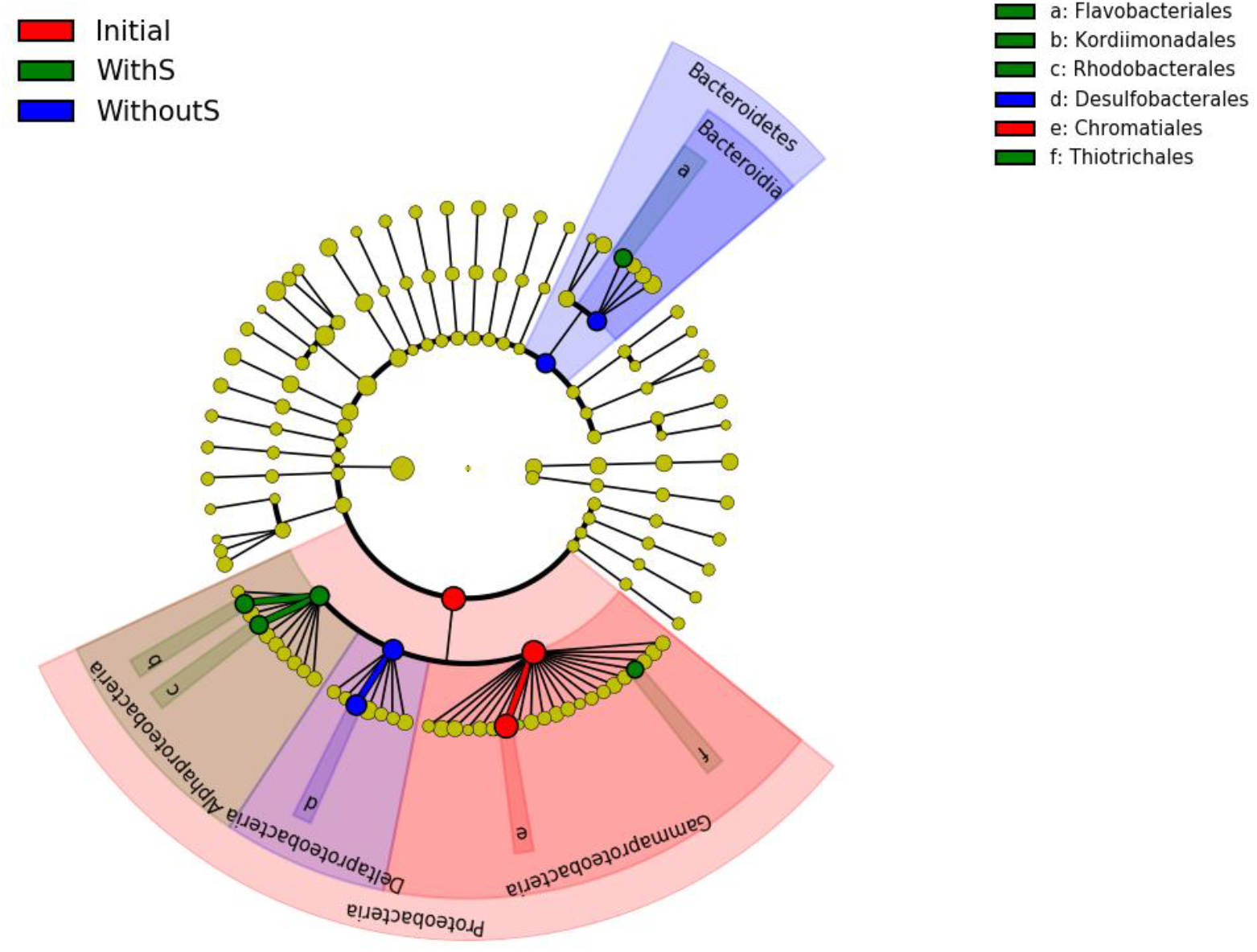
Linear discriminant analysis Effect Size (LEfSe) analysis (*p* < 0.05, LDA score > 2) of a class-level taxonomic cladogram.

The numbers of OTUs detected in the initial, with-S, and without-S groups were 272, 301, and 263, respectively (Fig. 3). According to the Venn diagram, 36 OTUs were shared among all experimental groups, 24 were shared between the initial and with-S groups, and 40 were shared between the reared groups regardless of sulfide presence (Fig. 3). The numbers of specific species in the initial, with-S, and without-S groups were 199, 201, and 174 OTUs, respectively.

**Figure 3.**
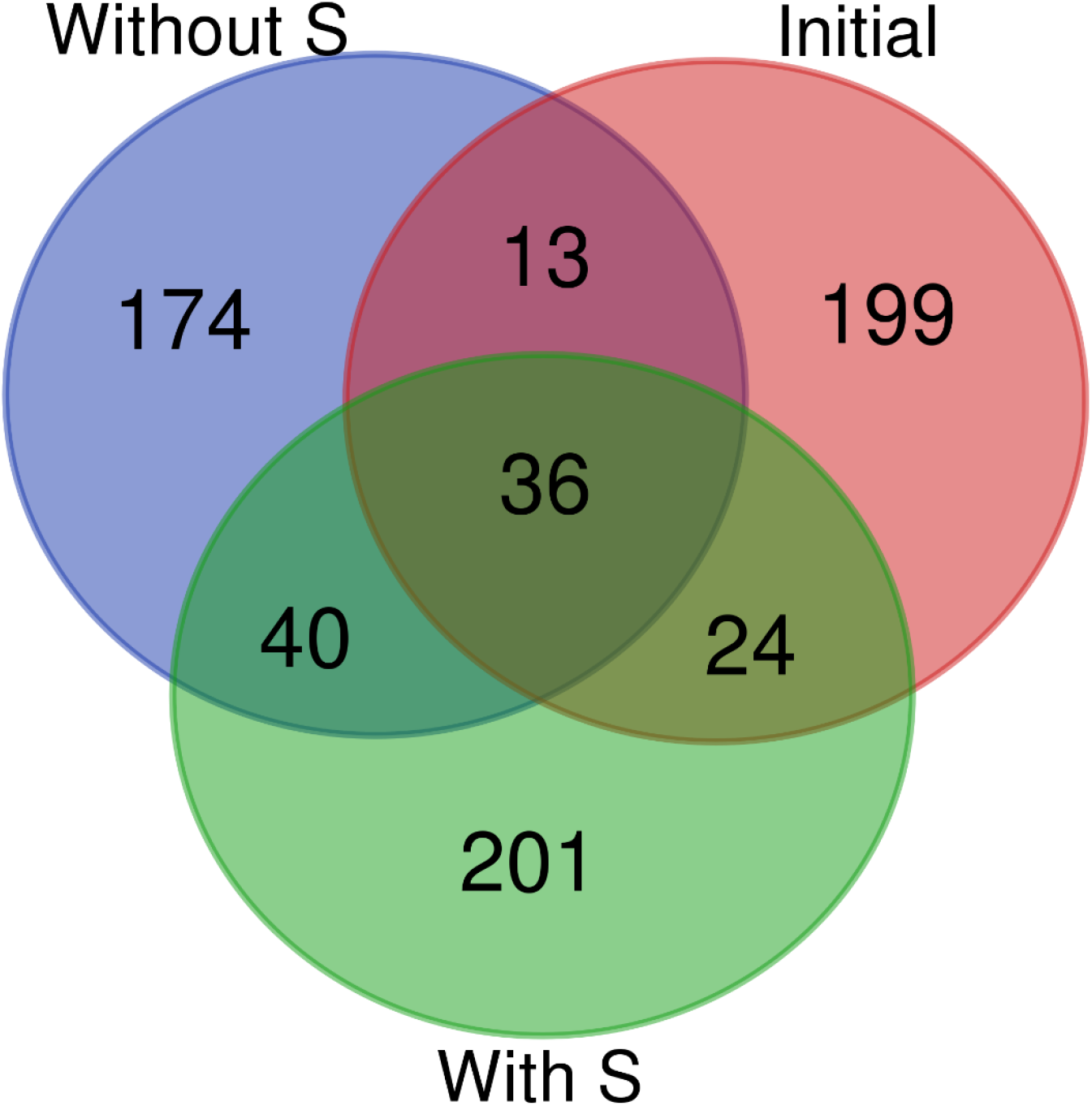
Venn diagram of OTUs detected in each experimental group. Numbers indicate the number of common and group-specific OTUs

### Abundance ratio and phylogenetic analysis

The sum of the counts of the top 10 OTUs accounted for more than 96% of total counts in all groups (Table 2). Several OTUs were shared among the experimental groups; specifically, two OTUs in the initial group, three in the with-S group, and four in the without-S group were also detected in the other experimental groups.

**Table 2.**
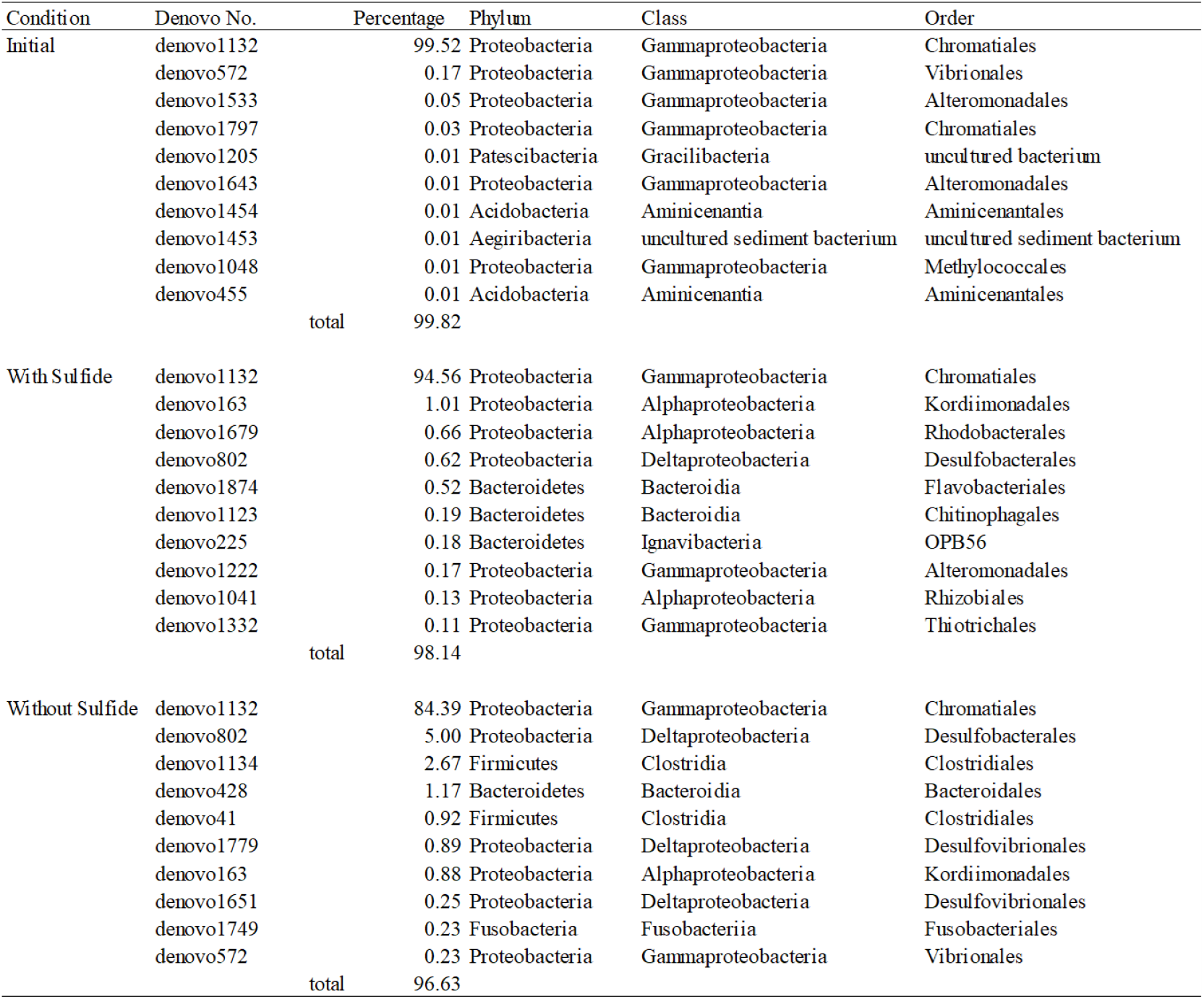
Top 10 OTUs ranked by abundance ratio in each experimental group. Species identification and OUT classification were performed using the SILVA database.

Denovo1132 was classified as γ-Proteobacteria and showed high counts in all experimental groups, which was consistent with previously reported sequences of symbionts of *L. satsuma* inhabiting Kagoshima Bay and the Nikko Seamount (Fig. 4). In contrast, Denovo1797, which was detected only in the initial sequence, differed by two bases from these sequences. Denovo572, Denovo1533, Denovo1643, Denovo1048, Denovo1222, and Denovo1332 were closely related to *Lamellibrachia* sp. symbionts in Sagami Bay and formed a distinct clade (Fig. 4). Paraphyletic members of the γ-Proteobacteria clade, Denovo1651, Denovo1779, and Denovo802, were detected in the reared groups and were classified as δ-Proteobacteria (Fig. 4). Denovo1454 and Denovo455 were detected only in the initial samples and were classified as Amnicenantia (Table 2). Denovo1134, Denovo41, and Denovo1749 were classified as Clostridia and Fusobacteria, respectively, and formed a sister group to α-Proteobacteria (Denovo1041, Denovo1679, and Denovo163) in this study (Fig. 4). The ε-Proteobacteria, previously reported as symbionts of *L. satsuma*, were relatively closely related to Denovo1205, which was detected in the initial samples. These bacteria were classified as Gracilibacteria. Denovo1453 was classified as Aegiribacteria and formed a clade with Denovo225 (Ignavibacteria) and Denovo428, Denovo1874, and Denovo1123 (Bacteroidia).

**Figure 4.**
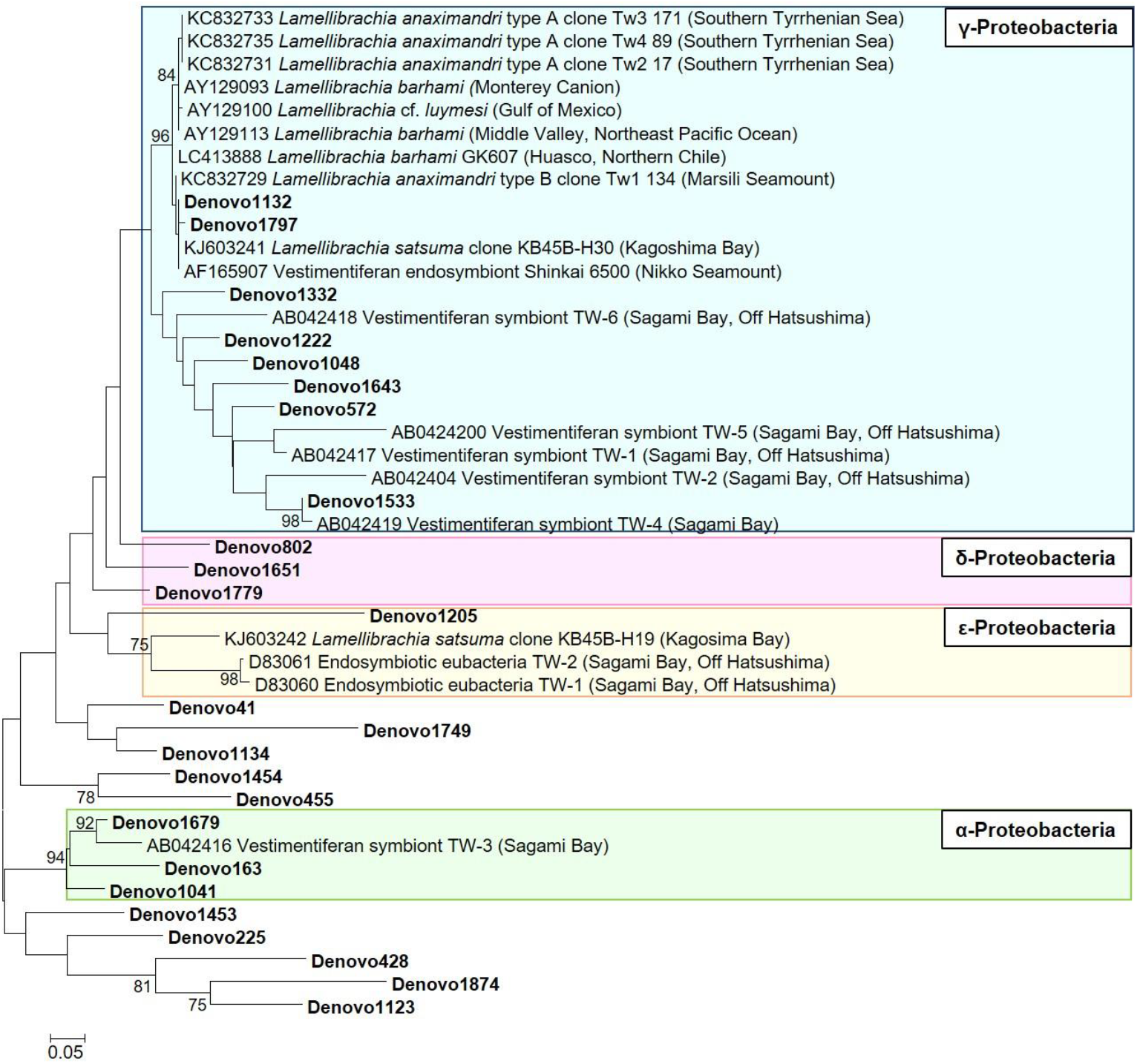
Maximum-likelihood phylogenetic tree constructed using the top 10 OTUs with the highest abundance ratios in each experimental group. Bootstrap values above 70 are shown. The abundance ratio of these OTUs are provided in Table 2.

**Figure 5.**
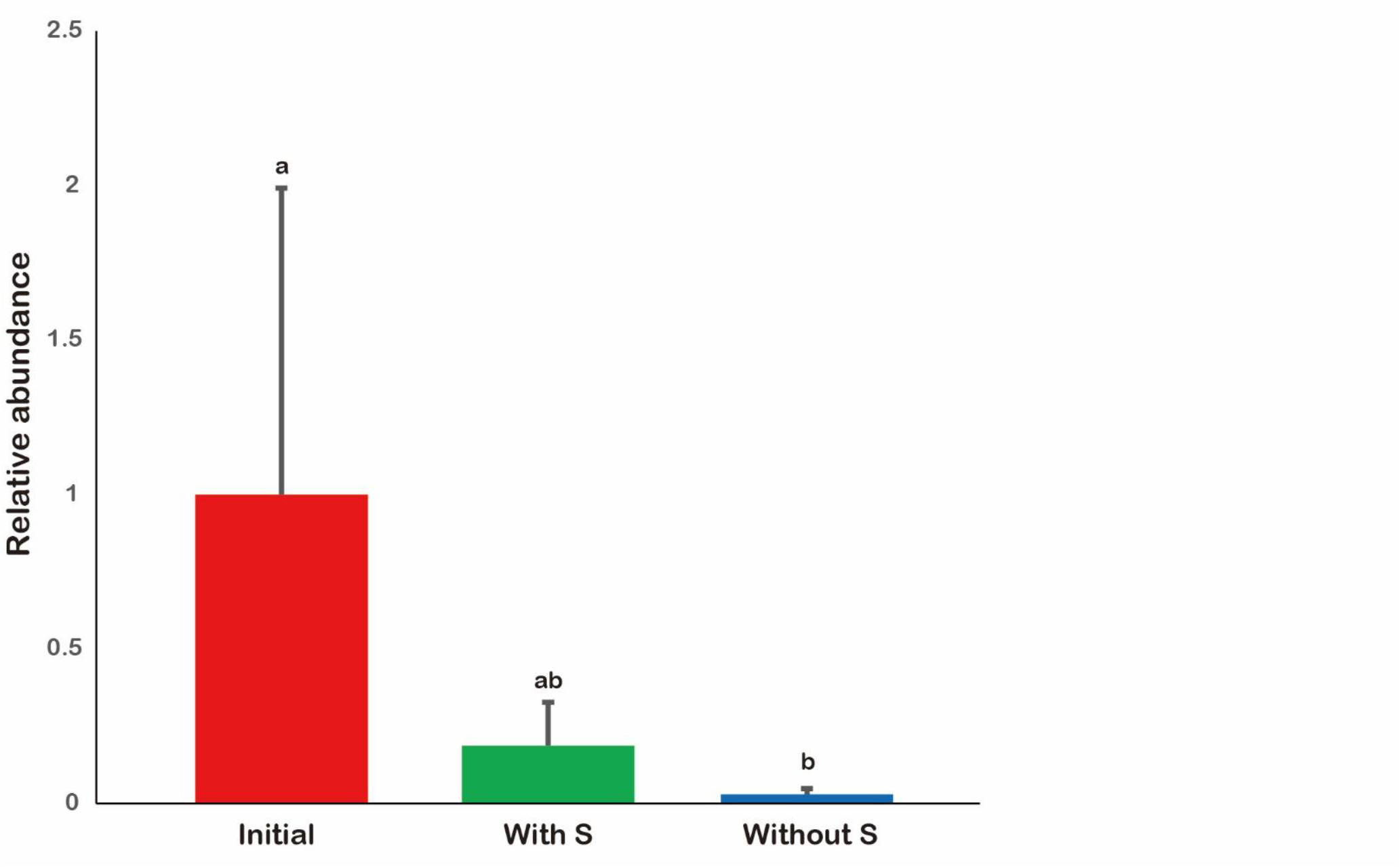
Relative quantification of the dominant γ-proteobacterial symbiont based on 16S rDNA levels in *Lamellibrachia satsuma* reared with or without sulfide supplementation. The mean values are shown relative to the initial levels upon collection (set to 1), with error bars representing standard deviation.

### Quantity of major symbionts

The results revealed that the with-S group did not differ significantly from the initial group, although a decreasing trend was observed. Conversely, the without-S group exhibited significantly different abundance.

## Discussion

This study provides the first qualitative insight into the highly diverse bacterial community within the trophosome of *L. satsuma* and reveals its dynamic changes under rearing conditions. Additionally, despite the significant decline in the abundance of the dominant symbiotic bacteria, the overall bacterial community did not exhibit a marked quantitative imbalance, and the host organism survived under both sulfide-supplemented and sulfide-free conditions.

The low microbial diversity observed immediately after collection is likely attributable to the dominance of a single major symbiont species and a lack of homogeneity with other microbial communities. In contrast, increased diversity under sulfide-free conditions was associated with the presence of numerous low-abundance species. This finding aligns with previous studies of the trophosomes of *Ridgeia piscesae* inhabiting the East Pacific Rise, which reported a lower diversity of microbial flora compared with the surrounding sediments and water, with Proteobacteria as the dominant phylum (He & Zhang, 2016). Similarly, *Bathymodiolus azoricus* collected from hydrothermal plume areas exhibited a marked reduction in symbiont populations under sulfide-free conditions, accompanied by an increase in non-symbiotic bacteria (Barros et al., 2018). Collectively, these findings indicate that the bacterial community within the trophosome of *L. satsuma* is less diverse in its natural habitat, with diversity shifting in response to rearing conditions.

From a quantitative perspective, the abundance of dominant symbiotic bacteria gradually declined under both sulfide-supplemented and non-supplemented conditions. Although sulfide supplementation did not maintain the original abundance of symbiotic bacteria, it slowed the rate of decline compared with non-supplemented conditions. These results suggest that while sulfide supplementation mimics natural conditions to some extent, it does not fully reproduce the complex interactions required to sustain symbiotic bacteria in their natural state. Similar observations have been reported for *B. azoricus*, in which symbionts were almost entirely lost within 30 days under sulfide-free environments but could be reacquired from the environment following sulfide reintroduction (Kádár et al., 2005). The relatively slow decline of symbiotic bacteria in *L. satsuma* is likely attributable to their low metabolic rate and slow growth, which are characteristics typical of tubeworms inhabiting cold seeps with limited sulfide availability (Fisher et al., 1997; Julian et al., 1999). Although Kagoshima Bay is classified as a hydrothermal vent system, *L. satsuma* has been suggested to utilize sub-seafloor, microbially derived hydrogen sulfide using methane in volcanic gases, a mechanism similar to that of cold-seep communities (Yamanaka et al., 2001).

This study also identified bacterial taxa that were either unique to or shared between experimental conditions. Common species across all conditions likely originated from the host’s natural habitat, whereas species shared between the reared groups may have resulted from rearing water colonization. Notably, α-Proteobacteria (*Kordiimonadales*) and δ-Proteobacteria (*Desulfobacterales*) were abundant under both rearing conditions but were absent immediately after collection, suggesting potential environmental uptake during rearing. These findings imply that adult *L. satsuma* may acquire bacteria through pathways such as gas exchange, reflecting the level of microbiome adaptability. The limited dominance of non-symbiotic bacteria under rearing conditions suggests that the trophosomal environment imposes constraints on microbial proliferation. Although contamination cannot be entirely ruled out for low-abundance species, differences in bacterial composition at the order level across conditions support the possibility of selective uptake or infection by the host. However, functional genes were not analyzed in this study, limiting our ability to draw definitive conclusions regarding the metabolic roles of these bacteria and their interactions with the host. Future studies incorporating functional gene analyses are needed to further clarify the ecological significance of these microbial dynamics.

*L. satsuma* was reared in two systems prepared by mixing natural and artificial seawater, with sodium sulfide solution added to one of the two systems. Consequently, no consistent pattern was observed when the same bacterial species were detected with or without sulfide supplementation. In particular, Proteobacteria and Bacteroidetes are present in diverse marine environments (Kirchman, 2002; Stevens et al., 2005). In contrast, the microbial microbiome in water has been reported to differ between shallow and deep waters (Sánchez-Soto Jiménez et al., 2018), and the microbiome of sediments and chimneys in areas of active or inactive hydrothermal activity, such as inactive chimneys, are rich in α-Proteobacteria and Nitrospirae (Sylvan; Toner & Edwards, 2012; Cerqueira et al., 2018; Hou et al., 2020). However, even if the bacterial species present in such environments overlap with those identified in the present study, their causal relationship with rearing conditions remains speculative.

Two strains of γ-Proteobacterial symbionts were identified. Tubeworm symbionts are often considered non-host-specific because of their horizontal transmission (Nelson and Fisher, 2000). There are cases in which different tubeworm species harbor the same symbiont (Brzechffa & Goffredi, 2021). Therefore, coevolution is not considered to occur, and the host habitat (hydrothermal vent or cold seep) or water depth has been suggested to define the symbiont species. In the present study, *Lamellibrachia* spp. from Sagami Bay and *L. satsuma* from Kagoshima Bay were not sympatric. Nevertheless, γ-Proteobacteria closely related to *Lamellibrachia* from Sagami Bay were detected despite their horizontal separation. In addition, the *Lamellibrachia* symbiont from Sagami Bay formed a clade distinct from the dominant symbiont (Denovo1132). A possible explanation is that the symbionts present when *Lamellibrachia* speciation occurred in the waters around Japan dispersed with the host. This is supported by the detection of α-Proteobacteria (Denovo1679) and ε-Proteobacteria (Denovo1205), which are closely related to *Lamellibrachia* from Sagami Bay.

Polymorphisms in symbiotic bacteria likely arose after the host dispersed and acquired a new niche. This may explain the detection of several γ-Proteobacteria differing by a few bases. During this cycle, tubeworm larvae acquire symbiotic bacteria that are released into the environment from dead individuals (Klose et al., 2015; Polzin et al., 2019). This suggests that habitats may dictate symbiont species and that symbiont species may reflect host speciation to some extent. However, 16S rRNA has been suggested to be unsuitable for detecting symbiont specificity and diversity because of its high sequence conservation (Breusing; Franke & Young, 2020). Furthermore, the sequences used in this study were short and the phylogenetic relationships were not robust. Therefore, further investigation is required.

## Conclusions

This study demonstrates that the symbiotic relationship in *L. satsuma* can withstand 6 months of rearing under both sulfide-supplemented and sulfide-free conditions, albeit with notable shifts in the bacterial community structure. Although the addition of sulfide slows the rate of decline of dominant symbiotic bacteria, it does not fully preserve the natural microbiome. Future research should focus on the functional roles of the identified bacterial taxa, their interactions with the host, and the long-term implications of the rearing conditions on the sustainability of these symbiotic relationships.

## Acknowledgements

We express our heartfelt gratitude to the crew of the Kagoshima University training vessel *Nanseimaru* and the staff of the Enoshima Aquarium for their invaluable support in conducting this study.

